# The metabolome and proteome of stem cell-derived human primordial germ cells: a multi-omics approach

**DOI:** 10.64898/2026.03.31.715517

**Authors:** Madalena Vaz Santos, Bauke V. Schomakers, Marta Llobet Ayala, Tara Jamali, Michel van Weeghel, Ans M. van Pelt, Callista L. Mulder, Geert Hamer

## Abstract

Primordial germ cells (PGCs) are the population of cells that, in the human embryo, specify day 12 post-fertilization, and form the precursor cells for the future egg or sperm cells. Although in vitro differentiation of PGCs from human stem cells has been achieved, these primordial germ cell-like cells (hPGCLCs) fail to further mature. The reason for this is unclear. Previous studies in mice revealed that several specific metabolic changes occur during the maturation of these cells, which are essential for their developmental progress. However, very little is known about the metabolic profile of human primordial germ cells. In the severe scarcity of human PGCs, hPGCLCs serve as a research model to study PGC formation.

To investigate this, we differentiated hPGCLCs using induced-pluripotent stem cells and performed a mass spectrometry analysis to establish their metabolome and proteome. These cells revealed distinct metabolic profile, with changes particularly at the proteome level. This included a shift between canonical and non-canonical citric acid cycle in hPGCLC, downregulation of late-stage glycolysis and reduction of nucleotide *de novo* synthesis. By providing an integrative map of these metabolic networks, we aim to provide insight on the influence of metabolism on human PGC development that could help improve methods for in vitro differentiation and maturation hPGCLCs.

## Introduction

Major breakthroughs have been achieved in recent years in the emerging field of in vitro gametogenesis (IVG), particularly in the generation of gametes from stem cells in the laboratory. In mice, both functional oocytes and spermatids have been generated from embryonic and induced pluripotent stem cells (Morohaku et al., 2016; Hayashi et al., 2017; Sun et al., 2018; Ishikura et al., 2021). These advances have sparked major practical and ethical discussions regarding possible clinical applications in the fields of fertility preservation and treatment (de Bruin et al., 2025).

However, in humans, using both embryonic stem cells and induced pluripotent stem cells (iPSCs), *in vitro* germ cell development has not progressed beyond the stage of human primordial germ cell-like cells (hPGCLCs). Primordial germ cells (PGCs) are the population of cells that arise in the human embryo around day 12 post-fertilization and later differentiate to form the foundation for gametogenesis (Chen et al., 2019; Castillo-Venzor et al., 2023). After the initial period of specification and along embryonic development, PGCs undergo a maturation process during which dynamic changes in their transcriptome occur alongside major rearrangements of their epigenetic landscape (Murase 2024, Guo 2015).

The restricted accessibility to human embryos limits the study of *in vivo* hPGC differentiation and highlights the importance of IVG research. In the past decades, various protocols for hPGCLC specification have been successfully developed, but it remains unclear why hPGCLCs fail to further mature (Irie et al., 2015; Overeem et al., 2023; Sasaki et al., 2015; Vijayakumar et al., 2023). To characterize the bonafide identity of the hPGCLCs, most studies focus on the analysis of the transcriptome and methylome (Vaz Santos et al., 2025). Although the transcriptome provides us with valuable information about the gene expression patterns, it does not offer a complete overview of the functional profile of the cell.

In the regulation of cellular function, metabolism plays a key role. Through its involvement in energy production, biosynthesis of essential macromolecules, and modulation of the epigenome, metabolism directly influences cellular differentiation and fate determination. (Verdikt et al., 2021, Haws et al., 2020, Hayashi et al., 2023, Tischler et al., 2019). Previous studies were able to show distinct metabolic features in regard to mouse PGC (mPGC) in comparison to their pluripotent stem cell progenitors (embryonic stem cells), while reporting how metabolism directly regulates germ cell development (Hayashi et al., 2018, Hayashi et al., 2024). It was shown that glycolysis decreases and oxphos increases along mPGC differentiation, with mouse PGCLCs (mPGCLCs) showing similar levels to the progenitors and early stage mPGCs (E9.5). Furthermore, inhibition of oxphos or glycolysis impaired early or late differentiation of mPGCLCs, respectively (Hayashi et al., 2018). Therefore, besides being essential for the development of working protocols for human IVG, other omics approaches, such as metabolomics and proteomics, are also required for mechanistic insight in the biology of hPGC formation and further differentiation of the human germ line. In addition to essential growth factors and a functional somatic niche (Lakshmipathi et al., 2025), both necessary to support continued germ cell development, it is possible that specific nutrients or oxygen/CO2 levels are also required to enable in vitro germ cell development beyond the PGC stage.

To enlight on the metabolic requirements of human germ cells, we performed a metabolomics and proteomics assessment, through mass spectrometry analysis comparing *in vitro* differentiated hPGCLCs, the non-hPGCLC fraction after differentiation and their progenitor human iPSCs. With this we detected shifts in the canonical and non-canonical Citric Acid cycle (TCA), alongside glycolytic pathway remodelling. This includes isoform switches and downregulation of late-stage glycolysis proteins. Additionally, the pentose phosphate pathway (PPP) also revealed changes, with indication of reduction of *de novo* nucleotide synthesis and increase in production of nucleoside precursors in hPGCLCs. By providing an integrative map of these metabolic networks, we intend to enlighten the influence of metabolism on the process of human PGC specification.

## Results

### Distinct metabolic and proteomic profiles between the progenitor, hPGCLC and non-hPGCLC groups

To investigate the metabolic profile and the changes that occur during the early stage of hPGCLC *in vitro* differentiation, we performed metabolomic and proteomic analyses using Ultra high performance liquid chromatography-mass spectrometry. For this, three groups of cells were analysed: (1) the iPSCs that served as the progenitors before in vitro differentiation; (2) the in vitro derived hPGCLCs, FACS sorted based on ITGA6+/EpCAM+ (3) the remaining ITGA6^-^ non-hPGCLCs. The hPGCLCs and non-hPGCLCs were sorted at the fifth day of differentiation and differentiation of iPSCs to hPGCLCs was confirmed using immunofluorescence staining for the PGC markers OCT4 and BLIMP1 (Figure S1). Principal component analysis (PCA) revealed that the metabolic and proteomic profiles of iPSCs were clearly distinct from those of the differentiated cells (Figure 1A). This suggests that most of the observed proteomic and metabolomic changes (Tables S1-S6) are associated with the general *in vitro* differentiation process of iPSCs, rather than specific differences between the differentiated hPGCLC and non-hPGCLC populations.

**Figure 1.**
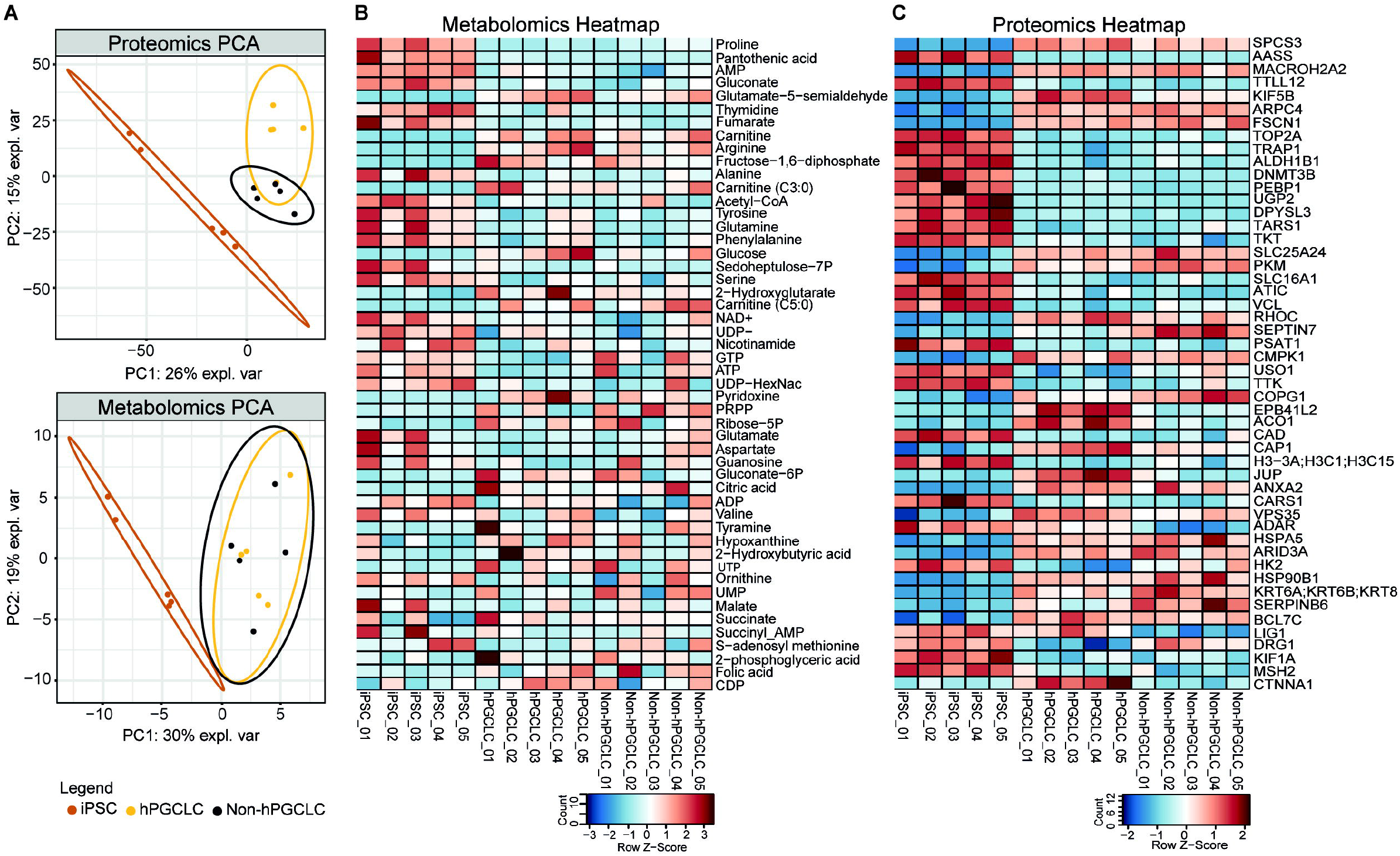
Multicomparison of iPSC, hPGCLC and non-hPGCLC groups. Five replicates were included per group for mass spectrometry analysis. **A**) PCA (principal component analysis) of the proteome and metabolome of the three groups. **B**) Heatmap of the top 50 metabolites and **C**) proteins, based on one-way Anova.

Regarding the proteome, all three groups clustered separately, with a clear separation between the progenitor cells (iPSCs) and differentiated cells (hPGCLCs and non-hPGCLCs) (Figure 1A). For the metabolome, PCA analysis showed that the hPGCLC and non-hPGCLC groups almost completely overlap, with only the progenitor cells grouping separately (Figure 1A). The top 50 metabolites and proteins that most significantly contributed to the difference between all groups are shown in the heatmaps (Figure 1B, 1C) and reflect the similarity between the differentiated groups (hPGCLC and non-hPGCLC) as well as the within-group variation at metabolite level.

The proteome analysis for the differentiated cell groups, hPGCLCs and non-hPGCLCs, identified a total of 487 significantly differentially expressed proteins (Table S4). Alongside proteins of the top 50 of most significant proteins (Figure 1C), the hPGCLCs showed upregulation of proteins associated with DNA repair mechanisms (LIG1, PARP1, ADAR), chromatin remodeling (CBX1, CBX3 and RBBP4), cell-cell adhesion (CTNNA1, CTNNA2, JUP, CTNNB1), nucleotide synthesis (PRPS1, PRPS2) and membrane transporters (SLC3A2, SLC2A3). On the other hand, hPGCLCs showed downregulation of proteins associated with cell cycle progression and division (CDK1, MAD2L1, SEPTIN7), translation factors and protein modification (EIF3F, EIF2S1, EIF3L, UBE2D2, UBE2D3), glycolysis metabolism (LDHA, LDHB) and retinoic acid signaling (CRABP2, RBP1).

Functional enrichment analysis of the differentially regulated proteins identified several enriched terms (Tables S7-S12). In comparison to the differentiated non-hPGCLCs, the hPGCLCs revealed enriched terms associated with cell adhesion and cytoskeleton, cellular traffic and localization, and indole-3-methanol response (Figure 2A). Simultaneously, the hPGCLCs showed a decrease of terms associated with nucleoside metabolism, sugar metabolism (namely Glycolysis/Gluconeogenesis) and central carbon metabolism (Figure 2A). In comparison with the progenitor iPSCs, the hPGCLCs showed enriched terms associated with cell adhesion and cytoskeleton, and cellular and organelle organization (Figure 2B). The downregulated terms were associated with translation, RNA binding and amino acid metabolism (Figure 2B). However, the comparison between non-hPGCLCs and iPSCs revealed similar up-and downregulated terms as those observed in comparison of the iPSCs with hPGCLCs (Table S7 and S8).

**Figure 2.**
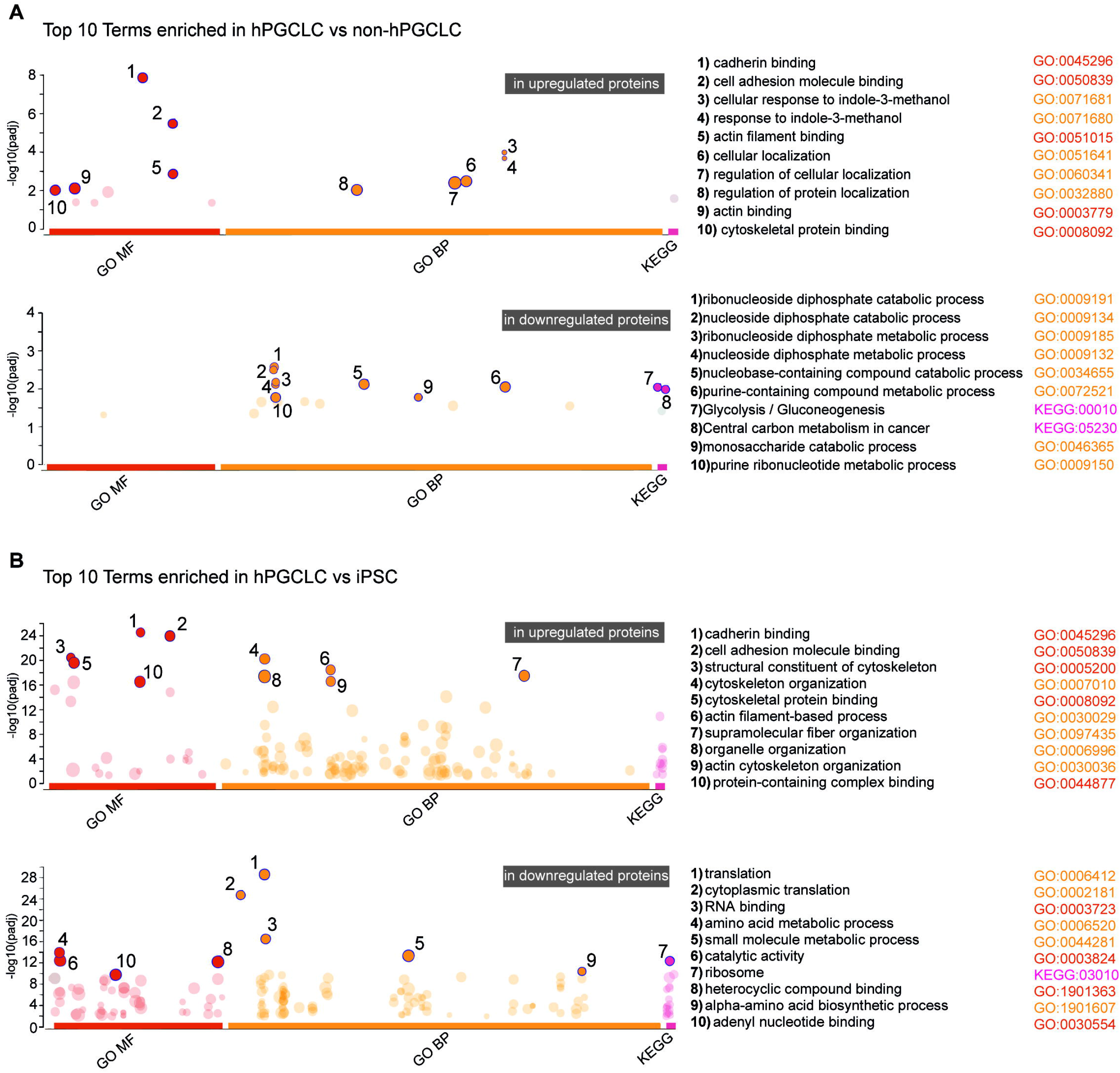
Top 10 enriched terms (based on p-value) of gene ontology (GO) analysis for significantly up-and downregulated proteins in **A**) hPGCLCs vs non-hPGCLCs and **B**) hPGCLCs vs iPSCs. Enrichment for terms in categories GO Molecular function (red), GO Biological process (orange) and KEGG pathways (magenta). Circles in transparent tone represent significantly enriched terms that are not part of the top 10; size of the circle represents number of proteins included.

### Energy metabolism of hPGCLCs is characterized by remodeling of the glycolytic pathway and upregulation of non-canonical TCA associated proteins

Energy production in the human cell relies mostly on two major pathways: glycolysis and oxidative phosphorylation (oxphos). Oxphos is preceded by the TCA, where the electron carriers required for the function of the electron transport chain (ETC) are produced.

Compared to iPSCs for energy-associated metabolites, hPGCLCs exhibited increased levels of glucose and fructose-1,6-diphosphate, and decreased levels of pantothenic acid, fumarate, ATP, AMP, and GTP (Figure 3A). No significant differences in metabolite levels were observed between the two differentiated cell types; both hPGCLCs and non-hPGCLCs showed similar changes relative to iPSCs, except that non-hPGCLCs did not display downregulation of ATP or GTP (Figure S2A).

**Figure 3.**
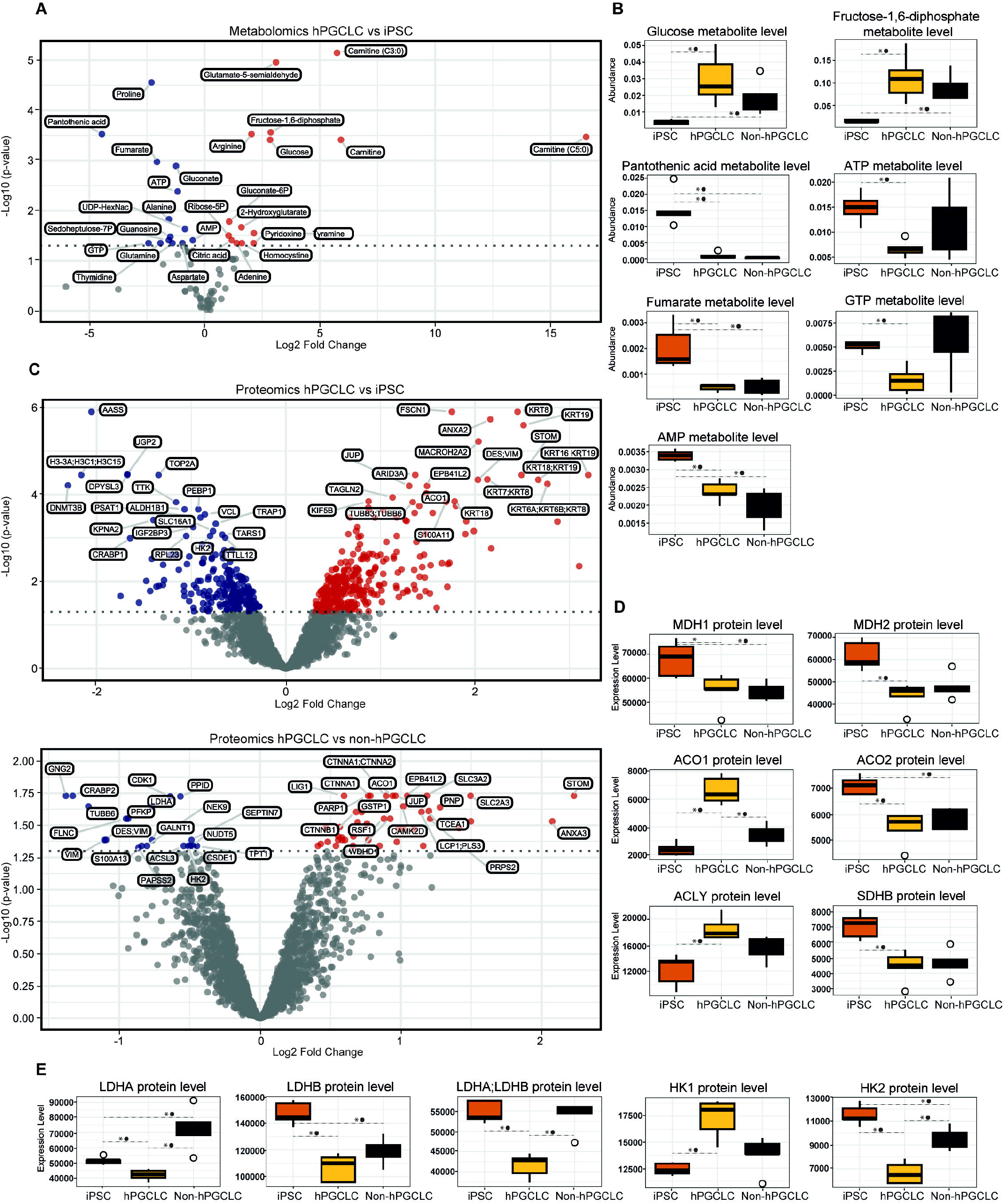
Energy metabolism of hPGCLCs is characterized by remodeling of the glycolytic pathway and upregulation of non-canonical TCA associated proteins. **A**) Volcano plot of the metabolome comparison between hPGCLCs *and* iPSCs; metabolites organized by p-value and increased in (red) or decreased in hPGCLCs (blue) abundance. **B**) Boxplot of metabolites related to glycolysis or oxidative phosphorylation. **C**) Volcano plot of the proteome comparison between hPGCLCs *and* iPSCs, and non-hPGCLCs *and* hPGCLCs; proteins organized by p-value and increased (red) or decreased (blue) expression. **D**) Boxplot of proteins related to oxidative phosphorylation and citric acid cycle or **E**) glycolysis. **\*** represents p value ≤ 0.05; **•** represents adjusted BH p value ≤ 0.05.

Overall, the proteome analysis revealed broad changes between all three groups associated to energy-related pathways, with up-and downregulation of proteins in both the TCA cycle and glycolysis (Figure 3C, D and E). For the TCA cycle, the hPGCLCs presented an upregulation of proteins constituent of the non-canonical TCA pathway, namely ACLY, ACO1 (*vs* iPSCs and non-PGCLCs) and IDH1 (*vs* non-PGCLC). Of the canonical TCA cycle, both differentiated cell types showed downregulation of the TCA associated proteins ACO2, SDHB, MDH1 and MDH2 in comparison to iPSCs (Figure 3C). For the ETC, although some proteins showed differences between cell types, no general pathway change was observed (Table S2, S4).

The glycolytic pathway also presented significant changes, with hPGCLCs revealing downregulation of enzymes associated with late-stage glycolysis, such as LDHA/LDHB (vs iPSCs and non-hPGCLC) and ENO1 (*vs* non-hPGCLC). Furthermore, a change in protein isoform levels was detected in hPGCLCs, with upregulation of HK1 in detriment of HK2 (vs iPSCs and non-hPGCLC) (Figure 3E).

An alternative energy source is through fatty acid oxidation. In this process, fatty acids are transported to mitochondria, via the carnitine shuttle, for subsequent beta-oxidation which will generate Acetyl-CoA, NADH and FADH2 (Carvalho et al., 2018). Both hPGCLCs and non-hPGCLC presented a high increase of acylcarnitine metabolites levels, in comparison to the progenitor cells (Figure 3A and S2A).

### hPGCLCs reveal heterogeneous changes in the PPP and indicate reduction of *de novo* nucleotide synthesis

The metabolic reprogramming of the cell encompasses more than energy production. A parallel pathway to glycolysis, the PPP, directly regulates biosynthesis capacity. This occurs due to its capacity to oxidize glucose, derived from glycolysis, to produce ribose-5-phosphate and NADPH, essential for redox balance and nucleotide synthesis, respectively (Kim et al., 2012).

At the metabolite level, compared to iPSCs, the hPGCLCs presented increased levels of ribose-5-phosphate and 6-phosphogluconate (activated form), while presenting decreased levels of gluconate (Figure 4A).

**Figure 4.**
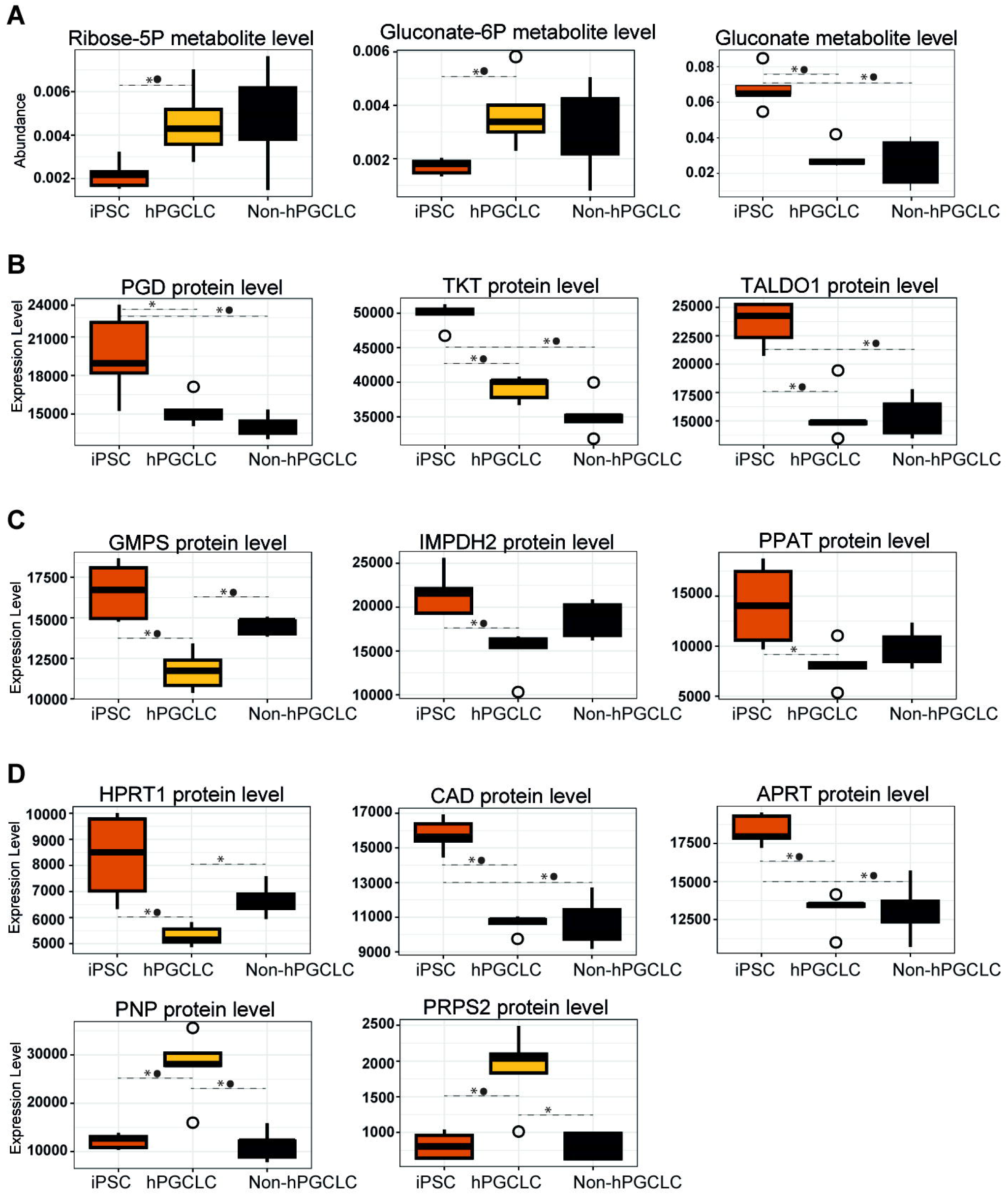
hPGCLCs reveal heterogeneous changes in the Pentose Phosphate Pathway (PPP) and indication of reduction of *de novo* nucleotide synthesis. **A**) Boxplot of metabolites involved in the Pentose Phosphate pathway (PPP) and nucleotide synthesis. Boxplot of proteins involved in **B**) PPP, **C**) *de novo* nucleotide synthesis and **D**) purine nucleotide salvage and nucleoside recycling. **\*** represents p value ≤ 0.05; **•** represents adjusted BH p value ≤ 0.05.

Similar to what was found for glycolysis and TCA pathways, multi-directional effects were detected in the PPP at the proteome level. As part of the oxidative PPP stage, the hPGCLCs presented downregulation of PGD (*vs* iPSCs). As part of the non-oxidative stage of the pathway, the hPGCLCs presented upregulation of TKT in comparison with the non-hPGCLCs and downregulation in comparison to the iPSCs. The protein TALDO, also part of the non-oxidative PPP, was downregulated (vs iPSCs and non-PGCLC) (Figure 4B).

Nucleotide synthesis is for a large part influenced by PPP activity, which provides the necessary components for this biosynthetic process (Kim et al. 2012). Related to this process, the hPGCLCs presented downregulation of GMPS and IMPDH2 (vs iPSCs and non-hPGCLCs), and PPAT and CAD (vs iPSCs), which are proteins associated to *de novo* purine and pyrimidine synthesis (Figure 4C). In addition, the hPGCLCs showed lower levels of HPRT1 (vs iPSCs and non-hPGCLCs) and APRT (vs iPSCs), related to purine salvage.

Furthermore, the hPGCLCs presented upregulation of PRPS2 and PNP (vs iPSCs and non-hPGCLCs), proteins associated with general purine synthesis substrate production and nucleoside degrading and recycling, respectively (Figure 4D).

## Discussion

The goal of this study was to analyze the metabolome and proteome of *in vitro-*derived hPGCLCs and how it differs from its progenitor iPSCs and parallelly differentiated non-hPGCLCs counterparts to better understand the energy metabolism of human PGCs. Overall, the hPGCLCs presented heterogeneous changes at the protein level in the PPP pathway, in comparison with both the iPSCs and non-hPGCLCs groups. Similarly, the nucleotide synthesis pathways revealed broad changes, suggesting that hPGCLCs opt for an energy cost-reduction of *de novo* synthesis while increasing nucleoside precursors.

The metabolite levels revealed a distinct profile between hPGCLCs and the progenitor cells, demonstrating that the metabolome changes with differentiation. Notably, the metabolome of hPGCLCs showed no significant differences from the parallelly differentiated non-hPGCLCs. However, the metabolic state is not always fully captured by metabolite levels at a single time point, as this approach does not account for changes in metabolic flux (Srivastava et al., 2016). The metabolite level might not change but the activity level, such as gain or loss of metabolites, might. In addition, some metabolites can be redirected differently. For example, Glucose-6-phosphate can be directed to either the PPP or glycolysis (Jiang Z et al., 2022). Nonetheless, a relevant finding is the high increase of acylcarnitine metabolites levels in both differentiated cell types in comparison to the progenitor cells. This could indicate that the observed alterations in the metabolome are a feature of differentiation in general from iPSCs. The increase of acylcarnitine metabolites could be indicative of alterations in fatty-acid oxidation, however no other findings in our dataset support this (Longo et al., 2016; Stephens et al., 2007). Acylcarnitine metabolites increase can also be indicative of cell-stress (Calabrese et al., 2006). As such, for the antioxidant effect, it could prove beneficial for the overall culture system to add acetyl-L-carnitine supplementation to the differentiation media.

The metabolic network not only depends on metabolites, but also on the abundance of regulatory proteins. Contrary to the metabolome, proteomic analysis revealed distinct metabolic profiles among all studied cell groups, suggesting that proteomic changes are a specific feature of differentiation to either hPGCLCs or non-hPGCLCs. It remains unclear why these proteomic changes do not correspond to differences in the metabolome. One possible explanation is that changes in protein expression may not have had sufficient time to result in measurable differences in metabolite levels. Overall, the results revealed distinct energy profiles between the hPGCLCs, non-hPGCLCs and the progenitor cells (iPSCs), characterized by significant changes at the proteome level in the TCA cycle, glycolytic pathway and pentose phosphate pathway.

In this study, the hPGCLCs presented upregulation of proteins constituent of the non-canonical TCA, while both hPGCLCs and non-hPGCLCs showed downregulation of proteins linked to the canonical TCA cycle. The shift between canonical and non-canonical TCA cycle has been previously described in multiple cell types (Mateska et al., 2022). In ESCs, the TCA cycle shifts from canonical to non-canonical when the cells differentiate and exit pluripotency and blocking the non-canonical TCA pathway was shown to impede pluripotency exit, showing its importance in general cell lineage specification (Arnold et al., 2022). The switch to the non-canonical pathway can redirect the TCA cycle to an anabolic purpose, increasing the biosynthetic capacity of the cell. Extensive epigenetic reprogramming is one of the main events that occurs during both *in vivo* PGC and *in vitro* hPGCLCs differentiation (Murase et al., 2024; Sugawa et al., 2025). It is known that the switch to the non-canonical pathway can lead to increased production of acetyl-CoA and regeneration of NAD+, which can influence epigenetic reprogramming through histone acetylation and promote glycolysis, respectively (Doan et al., 2022; Arnold et al., 2022). While our data indicates upregulation of proteins constituent of the non-canonical TCA in hPGCLCs, no increase of NAD+ or acetyl-CoA was observed. In addition, no alpha-ketoglutarate was detected in our analysis, which was previously described as a key metabolite connecting metabolism and epigenetic reprogramming in mouse germ cells (Xing et al., 2020; Tischler et al., 2019). Overall, the observed changes could indicate metabolic regulatory rewiring, leading to a preference for non-canonical TCA in hPGCLCs. Furthermore, the functional enrichment analysis identified glycolysis/gluconeogenesis as a top ten downregulated term in hPGCLCs in comparison to non-hPGCLCs. Together with the downregulation of late-stage glycolysis associated proteins, this could suggest a metabolic reprogramming that directs the hPGCLCs to a less proliferative, more biosynthesis prone state.

In addition to the late-stage glycolysis, our data suggests further remodelling of the glycolytic pathway in hPGCLCs. We observed that these cells show a switch in isoform preference between the enzymes of the first-committing step in glycolysis, in comparison to both iPSCs and non-hPGCLCs, from HK 2 (predominantly active in iPSC) to HK1. Previous research, in mice, has associated HK2 prevalence with highly glycolytic cells, which favour proliferation, such as cancer or stem cells (Wolf et al., 2011). HK1, on the other hand, can act to promote the shift of glucose-6-phosphate from the mitochondria to the cytoplasm, with the potential for metabolic rerouting of this substrate between glycolysis and PPP pathways (De Jesus et al., 2023). Based on these results, the efficiency of glycolytic inhibitors for use on hPGCLCs, such as 2-Deoxy-D-glucose (commonly used in essays like the Seahorse Extracellular Flux) should be evaluated. Although this inhibitor acts by inhibiting the hexokinases, some studies suggest that it predominately affects HK2 function (Tseng et al., 2018; Wang et al., 2014; Tsai HJ, Wilson JE. 1996). Considering the HK1 isoform preference in hPGCLCs, the use of this inhibitor in this cell type might not be adequate.

The pentose phosphate pathway (PPP) contributes to nucleoside production, biosynthetic capacity and redox control of the cell, through production of ribose-5-phosphate and NADPH. For its function this pathway requires the use of the substrate glucose-6-phosphate, produced in the early steps of the glycolytic pathway (Jiang et al., 2014). Our data suggests modulation of PPP activity in hPGCLCs rather than a general activation or suppression, as evidenced by alterations on both the oxidative and non-oxidative stages of this pathway. Particularly, the changes in 6-Phosphogluconate dehydrogenase (PGD) levels could suggest modulation of the NADPH production rate to maintain redox homeostasis (Dubreuil et al., 2020; Ju et al., 2020). Combined with the observed increase in metabolite levels, this could indicate an increase of the oxidative PPP flux and possible biosynthetic capacity, when comparing to the progenitor cells (Jiang et al., 2014; Polat et al., 2021). However, since no changes in NADPH or glutathione were detected, which are central predictors of the antioxidant and redox capacity of the cell, there is no direct evidence for alterations in redox metabolism. When comparing to their differentiated counterpart (non-hPGCLCs), the increased levels of TKT could suggest a higher metabolic flexibility of the hPGCLCs, through interconversion of glycolytic products and pentose phosphates (Kim et al., 2012; Chen et al., 2022).

Nucleotide synthesis depends on ribose-5-phosphate derived from the PPP (Jiang et al. 2014). The hPGCLCs presented downregulation of proteins associated with de novo synthesis, while showing upregulation of proteins related to nucleoside recycling and nucleotide substrate production, in comparison to the iPSCs and non-hPGCLCs. Furthermore, the functional enrichment analysis revealed a downregulation of terms associated with nucleoside metabolism, in comparison to non-hPGCLCs. This pattern could indicate a shift in the germ cell metabolic function by maintaining the nucleotide pool through less energy-demanding mechanisms, a characteristic of less proliferative, specialized cells (Madsen et al., 2023; Lane et al., 2015). Such metabolic restraint would align with the suggested energy-quiescent profile of hPGCLCs, enabling them to allocate resources toward biosynthetic processes and genomic stability.

Similar research has been done regarding metabolism in mouse PGCs (mPGCs). The metabolic profile of mPGCs revealed a transition from a glycolytic state at (E9.5), similar to their ESCs progenitors, to a mainly oxidative state (E13.5) (Hayashi et al. 2017). While our hPGCLCs presented pinpoint pathway changes instead of overall pathway transitions, the observed alterations targeted several of the same pathways as in mice. Both the hPGCLCs and mPGCs demonstrated changes at the TCA level: in the mouse system particularly in alpha-ketoglutarate regulation (Hayashi et al., 2017; Tischler et al., 2018), in our human system more focused on the non-canonical TCA. A similar parallel can also be drawn at the glycolysis level, with the mPGCs shifting from a glycolytic precursor to a highly oxphos state (Hayashi et al. 2017), while the protein changes in hPGCLCs suggest downregulation of late-stage glycolysis. Furthermore, the changes observed in this study at the nucleotide synthesis level also relate to what was previously described in mouse cells (Hayashi et al., 2017), indicating that both models seem to tend towards an energy-efficient nucleotide pool maintenance. When looking at the *in vitro* differentiated mPGCLCs, the energy metabolism of these cells was more similar to that of their progenitor cells rather than mPGCs. In addition, the extracellular acidification rate and oxygen consumption rate of mPGCLCs indicated lower oxphos and similar glycolytic activity levels in comparison to mPGCs, suggesting that these levels are altered at a later stage of differentiation (Hayashi et al., 2017). Such findings align with the results shown here for hPGCLCs, with rearrangements of components of both the TCA and glycolysis pathways. Overall, this could indicate that the maturation status of germ cells is an important factor for their metabolic rewiring since mPGCLCs correlate to an earlier developmental stage than the mPGCs and closely resemble the energy profile of their ESC progenitors. For further understanding of the evolving metabolic profile in hPGCLCs, it would be helpful to include *in vitro* differentiated samples that correspond to more mature hPGCLC developmental stage, allowing for a better mapping of the metabolic development along human germ cell differentiation.

Overall, the data presented in this study indicate that *in vitro* differentiation of iPSCs to hPGCLCs is characterized by a metabolic regulatory rewiring, mainly at the protein level. This is reflected by a more quiescent cell state, with modulation of specific pathways that include a preference for non-canonical TCA activity, downregulation of late-stage glycolysis regulators and specific protein isoform switches in energy-related pathways. However, although these changes were shown at protein level, they did not translate to metabolome changes. This was previously also observed in mPGCLCs and suggests the importance of further maturation for complete metabolic rewiring to take place (Hayashi et al., 2017). Nevertheless, the newly obtained knowledge provides a better understanding of the initial steps of the metabolic reprogramming during early human germ cell development.

## Methods

### Human iPSC culture

The human iPSCs (LUMC0054iCTRL03 male cell line; LUMC iPSC core facility) (Overeem et al., 2023) were maintained in culture at 37C, 5% CO2 with use of mTESR plus media, for a maximum number of thirty-five passages. Every 5-7 days, the cells were passaged, with the use of ReLSR, to a Geltrex coated well containing mTESR plus media, 1% PenStrep and 1% RevitaCell. The day after passaging, the media was refreshed for removal of RevitaCell. The media was refreshed every other day. Mycoplasma testing was performed monthly on cultured iPSCs. The iPSCs were kept in culture for no more than 15 passages, to a maximum passage number of 30. Frozen iPSC stocks were stored in liquid nitrogen. RNA expression of pluripotency markers *Oct4, Nanog* and *Sox2* in undifferentiated iPSCs was confirmed by quantitative (q) PCR (Figure S3). qPCR data was analyzed with RDML GEAR Tools v1.4 (Untergasser et al., 2026).

### hPGCLCs differentiation and isolation

hPGCLCs were differentiated according to a previously published protocol (Overeem et al., 2023). At day four of differentiation, the heterogeneous population was prepared for fluorescence-activated cell sorting (FACS). For this, the differentiated cell population was dissociated into a single-cell suspension upon incubation for 10 min with Accutase. After washing, the cells were incubated with a live-dead marker (Zombie-green, Biolegend, cat. 423112) at a concentration of 0.5uL per million cells, for 30min at room temperature. The cells were then washed twice, for 5mins, with FACS buffer (PBS with 0.5% BSA) and incubated with the directly conjugated antibodies BV421-ITGA6 (Biolegend, cat. 313624) and APC-EpCAM (Biolegend, cat. 324208) at a concentration of 5uL per million cells, for 30min at 4C. After washing, the cell pellet was reconstituted in FACS buffer and the cell sorting was performed using a Sony SH800 sorter machine. Both ITGA6+/EpCAM+ (hPGCLCs) and ITGA6-/EpCAM-(Non-hPGCLCs) cell populations were sorted and collected for posterior analysis. In parallel, the progenitor iPSC population used to initiate the differentiation trial was also retrieved, through dissociation of the colonies with Accutase and collection of the pellet. Upon collection of the cell samples, dry pellets corresponding to 250.000 cells per sample were snap freeze and stored at –80C until further processing.

### Mass spectrometry and data analysis

A total of five replicates, per group (iPSCs, hPGCLCs and non-hPGCLCs), collected from five independent differentiation trials, were included for mass spectrometry analysis. The metabolomics and proteomics unified extraction was performed combining previously described protocols (Schomakers BV, et al. 2022; Schomakers BV, et al. 2025; Szyrwiel L, et al. 2024; Szyrwiel L, et al. 2023; Cox J, et al. 2014). The metabolites were analyzed with a Waters Acquity UHPLC together with a Bruker Impact II™ Ultra-High Resolution Qq-Time-Of-Flight mass spectrometer and data analysed using Bruker TASQ software version 2.1.22.3. Metabolites were normalized for total metabolite abundance. The proteomics sample preparation was done with a Thermo Scientific™ EasyPep™ MS Sample Prep Kit (A40006) and proteins were analysed with a Waters™ Acquity UPLC paired with a Bruker timsTOF Pro 2 mass spectrometer. Protein levels were normalized using method MaxLFQ (Cox J, et al. 2014). Statistical analysis and (adjusted) p value calculations were performed with R Studio package Limma 3.64.3.

Functional enrichment (Gene Ontology) analysis was performed with the differentially expressed proteins, up-and downregulated in separate, using g:Profiler g:GOSt (Kolberg et al. 2023). Protein identifiers were mapped with the background of all annotated human genes, with g:SCS multiple test correction and a threshold for significance of 0.05. Result terms were obtained for Gene Ontology Biological Process, Gene Ontology Molecular Function and KEGG Pathway.

### Immunofluorescence

For immunofluorescence staining, the cells were cultured in a Lab-Tek chamber slide with removable wells and fixed, with 4% PFA for 10 minutes, at day 4 of hPGCLCs differentiation. After three washes with PBS, the cells were incubated with 0.5% Triton/PBS for 15 minutes, at room temperature. Following three washes with 0.1% PBST, the cells were incubated with Superblock for one hour, at room temperature. Subsequently, the Superblock was removed and the cells were incubated overnight, at 4C, with the primary antibodies Oct3/4 (santa cruz, mouse sc-5279; 1:200) and Blimp1 (invitrogen, rat 14-5963-82; 1:100), diluted in Brightdiluent. In parallel, for negative controls, one well was incubated with mouse Igg and rat Igg in the same concentrations as the primary antibodies. The following day, the cells were washed three times with 0.1% PBST. Next, they were incubated, for 1h at room temperature, with the secondary antibodies anti-mouse Alexa Fluor 488 (1:1000) and anti-rat Alexa Fluor 555 (1:1000), and DAPI (1:1000). Lastly, the cells were washed three times with PBS and mounted with ProLong Gold.

More details on methodology are described in supplemental methods.

## Acknowledgements

The funding for this work was provided by ZonMW PSIDER (Program No. 10250022120001) (HipGametes). We thank Susana M. Chuva de Sousa Lopes for sharing the human iPSCs within the HipGametes consortium.

