## Supplementary figures and information for "The metabolome and proteome of stem cell-derived human primordial germ cells: a multi-omics approach"

### Supplements

#### Supplemental methods

##### Sample processing, mass spectrometry and data analysis

The metabolomics and proteomics unified extraction was performed combining previously described protocols (Schomakers BV, et al. 2022; Szyrwiel L, et al. 2024; Szyrwiel L, et al. 2023; Cox J, et al. 2014). For metabolomics, the following quantities of internal standard dissolved in water, were added to each sample tube containing 250.000 cells in a dry pellet: Adenosine-15N5-monophosphate (5 nmol), Adenosine-15N5-triphosphate (5 nmol), Alanine-13C3,15N (5 nmol), Arginine-13C6 (5 nmol), Aspartic acid-13C4 (5 nmol), D3-Carnitine (0.5 nmol), D4-Citric acid (0.5 nmol), 13C1-Citrulline (0.5 nmol), Cystine-13C6,15N2 (2.5 nmol), 13C6-Fructose-1,6-diphosphate (1 nmol), Glutamic acid-13C5 (5 nmol), D5-Glutamine (0.5 nmol), Glycine-13C2,15N (5 nmol), Guanosine-15N5-monophosphate (5 nmol), Guanosine-15N5-triphosphate (5 nmol), 13C6-Glucose (10 nmol), 13C6-Glucose-6-phosphate (1 nmol), Histidine-13C6 (5 nmol), Isoleucine-13C6, 15N (5 nmol), D3-Lactic acid (1 nmol), Leucine-13C6,15N (5 nmol), Lysine13C6 (5 nmol), Methionine13C5,15N (5 nmol), D6-Ornithine (0.5 nmol), Phenyl-13C6-alanine (5 nmol), Proline-13C5 (5 nmol), 13C3-Pyruvate (0.5 nmol), Serine-13C3,15N (5 nmol), D6-Succinic acid (0.5 nmol), Threonine-13C4 (5 nmol), D4-Thymine (1 nmol), Tyrosine-(phenyl-13C6) (5 nmol), D5-Tryptophan (0.5 nmol), Valine-13C5 (5 nmol).

In the same tube, lipid layer separation was performed (not included in this study). After, solvents were added until reaching a volume of 500uL water, 500uL methanol. Sequentially, 1mL of chloroform was added to each tube and samples were properly mixed and then centrifuged for 10min at 14.000 rpm for layer separation.

The top layer, consisting of the polar phase, was moved to a clean tube and dried with resort to a vacuum concentrator. Posteriorly, the dried samples were reconstituted in 100uL 6:4 methanol/water and metabolites were analyzed with a Waters Acquity UHPLC together with a Bruker Impact II™ Ultra-High Resolution Qq-Time-Of-Flight mass spectrometer. The chromatography column used for separation was a Merck Millipore SeQuant ZIC-cHILIC column, kept at 30 °C. Mass spectrometry (MS) data was acquired using negative and positive ionization in full scan mode over the range of m/z 50-1200, and analysed using Bruker TASQ software version 2.1.22.3. All reported metabolite intensities were normalized to freeze-dried cell number, as well as to internal standards with comparable retention times and response in the MS.

Metabolite identification was based on a combination of accurate mass, (relative) retention times, ion mobility data and fragmentation spectra, compared to the analysis of a library of standards. Metabolite quantification was done with ratio of a selected metabolite against the internal standards.

After removal of lipid layer, the protein pellet was dried using nitrogen and the sample was prepared using a Thermo Scientific™ EasyPep™ MS Sample Prep Kit. Samples were reduced and alkylated at 95°C for ten minutes, followed by a two-hour incubation at 37°C with a Trypsin/Lys-C protease mixture. Posteriorly to sample clean-up with the Peptide Clean-up Plate, samples were dried with nitrogen and sequentially resuspended in a 100 µL mixture of 97:3 (v/v) water:acetonitrile, containing 0.1% formic acid. Proteins were analysed with a Waters™ Acquity UPLC, using a Waters™ Acquity UPLC BEH C18 chromatography column, kept at 60 °C. MS data was acquired with a Bruker timsTOF Pro 2, using positive ionization in DIA-PASEF mode as previously reported (Szyrwiel L, et al. 2023). DIA-PASEF files were analysed with DIA-NN version 1.8.1 (Demichev et al., 2020), using a spectral library provided by Bruker. Data were normalized in DIA-NN using MaxLFQ.

#### Quantitative (q)PCR

Three distinct passages (p22, p23 and p27) of independent iPSC cultured samples were used for this analysis. RNA was isolated with a RNeasy Plus Mini Kit according to the manufacturer's instructions (Qiagen, cat. 74134). cDNA was synthesized with M-MLV Reverse transcriptase (RT) kit (Promega, cat. M1705). Mastermix I contained, per sample, 1ug RNA, 1ug of 500ng/uL Random Primers (Invitrogen, cat. 48190-011) and H2O up to a total volume of 15uL. Upon vortex, the mix was heated to 70C for 5 minutes. Master mix II contained, per sample, 5uL of 5x First Strand Buffer, 1.25uL of 10mM dNTP mix (Thermofisher, cat. R0192), 0.625uL RNasin 40U/uL (Promega, cat. 2511), 1uL of 200U/uL M-MLV RT and H2O up to a total volume of 10uL. Upon combining 15uL of master mix I with 10uL of master mix II, the sample was incubated at 25C for 10 minutes, followed by 1hour at 37C and forever at 4C. Minus RT control was produced simultaneously. Quantitative PCR analysis was performed with a LightCycler 480 (Roche), with SYBR Green Mix (Roche, cat. 4887352001). Primers were used at a final concentration of 0.5uM, for target genes' *Oct4* (Fw:AGGAATCGGGCCGGGGGTTG/ Rv:GTCCTGGGACTCCTCCGGGT; annealing temperature 68C), *Nanog* (Fw:CCTGTGATTTGTGGGCCTGA/ Rv:GGGTTGTTTGCCTTTGGGAC; annealing temperature 68C) and *Sox2* (Fw:GTTACGCGCACATGAACGG/ Rv:GTAGGACATGCTGTAGGTGGG; annealing temperature 65C), and for reference genes' *ActB* (Fw: CCTCGCCTTTGCCGATCC/ Rv: CCACCATCACGCCCTGG; annealing temperature 62C) and *Tmed1* (Fw: GGCCTGGAAGACCACCT/ Rv: GCCACACCACCCTTTGTT; annealing temperature 62C). In addition to three technical replicates (wells), minus RT and blank controls were included. qPCR data was analysed with RDML GEAR Tools v1.4 (Untergasser et al., 2026).

#### Supplemental Tables

Table S1. Metabolite counts and statistical analysis for the comparison hPGCLC vs iPSC

Table S2. Protein counts and statistical analysis for the comparison hPGCLC vs iPSC

Table S3. Metabolite counts and statistical analysis for the comparison hPGCLC vs non-hPGCLC

Table S4. Protein counts and statistical analysis for the comparison hPGCLC vs non-hPGCLC

Table S5. Metabolite counts and statistical analysis for the comparison iPSC vs non-hPGCLC

Table S6. Protein counts and statistical analysis for the comparison iPSC vs non-hPGCLC

Table S7. Functional enrichment analysis results for upregulated proteins in comparison hPGCLC vs iPSC

Table S8. Functional enrichment analysis results for downregulated proteins in comparison hPGCLC vs iPSC

Table S9. Functional enrichment analysis results for upregulated proteins in comparison hPGCLC vs non-hPGCLC

Table S10. Functional enrichment analysis results for downregulated proteins in comparison hPGCLC vs non-hPGCLC

Table S11. Functional enrichment analysis results for upregulated proteins in comparison non-hPGCLC vs iPSC

Table S12. Functional enrichment analysis results for downregulated proteins in comparison non-hPGCLC vs iPSC

#### Supplemental Figures

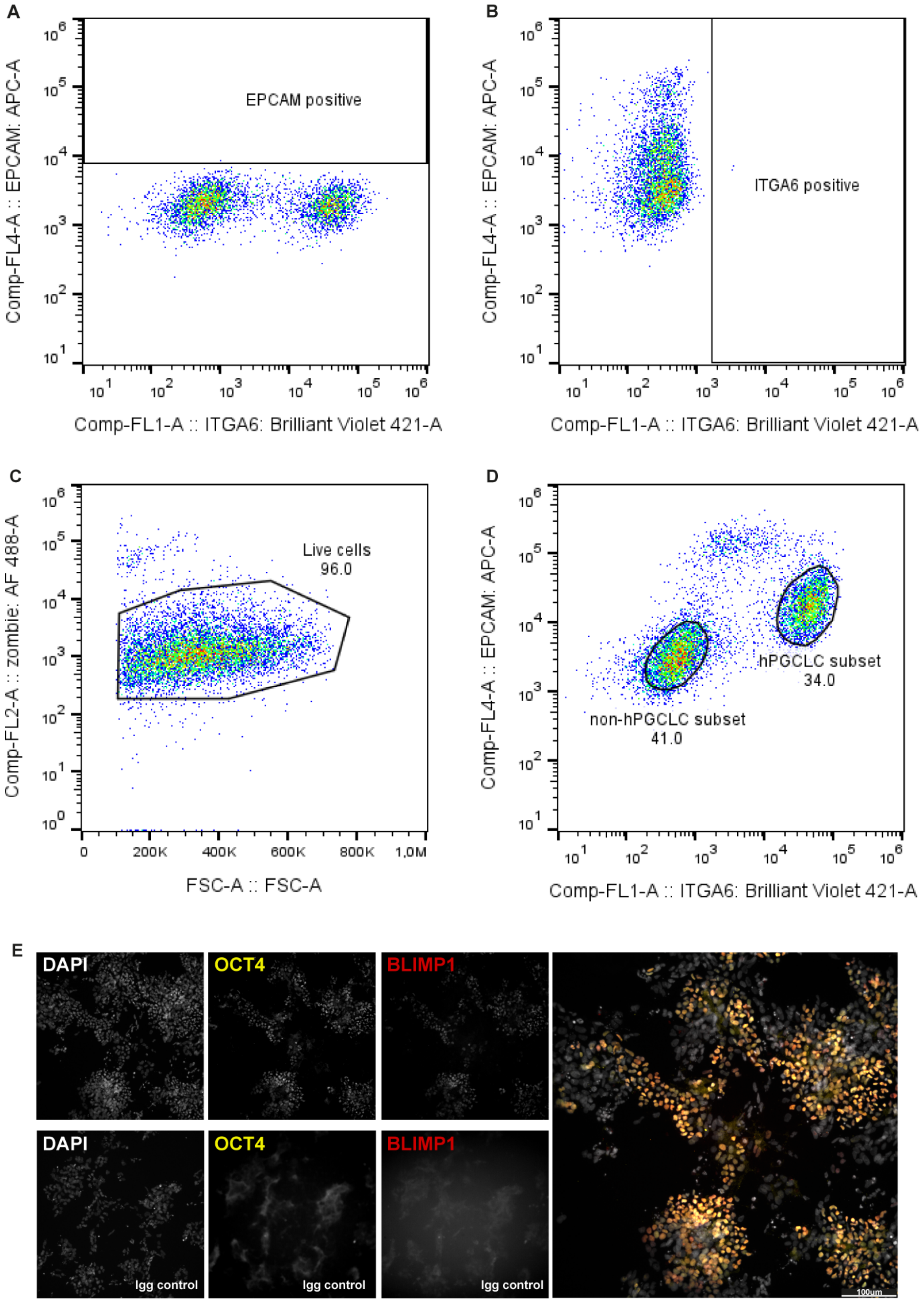

**Figure S1** – Validation of hPGCLC differentiation through immunofluorescence and Fluorescence-activated cell sorting of hPGCLC and non-hPGCLC groups, based on antibodies ITGA6 (biolegend, 313624) and EpCAM (biolegend, 324208) staining. **A**) FACS Fluorescence minus-one EpCAM control, **B**) FACS Fluorescence minus-one ITGA6 control, **C)** FACS Live cell population staining (Zombie AF 488) and selection, and **D)** FACS Gating of hPGCLC and Non-hPGCLC populations for sorting; **E)** Immunofluorescence staining of pre-sorting hPGCLC (and non-hPGCLC) population at day 4 of differentiation, for DAPI and primary antibodies Oct3/4 (santa cruz, sc-5279; 1:200) and Blimp1 (invitrogen, 14-5963-82; 1:100).

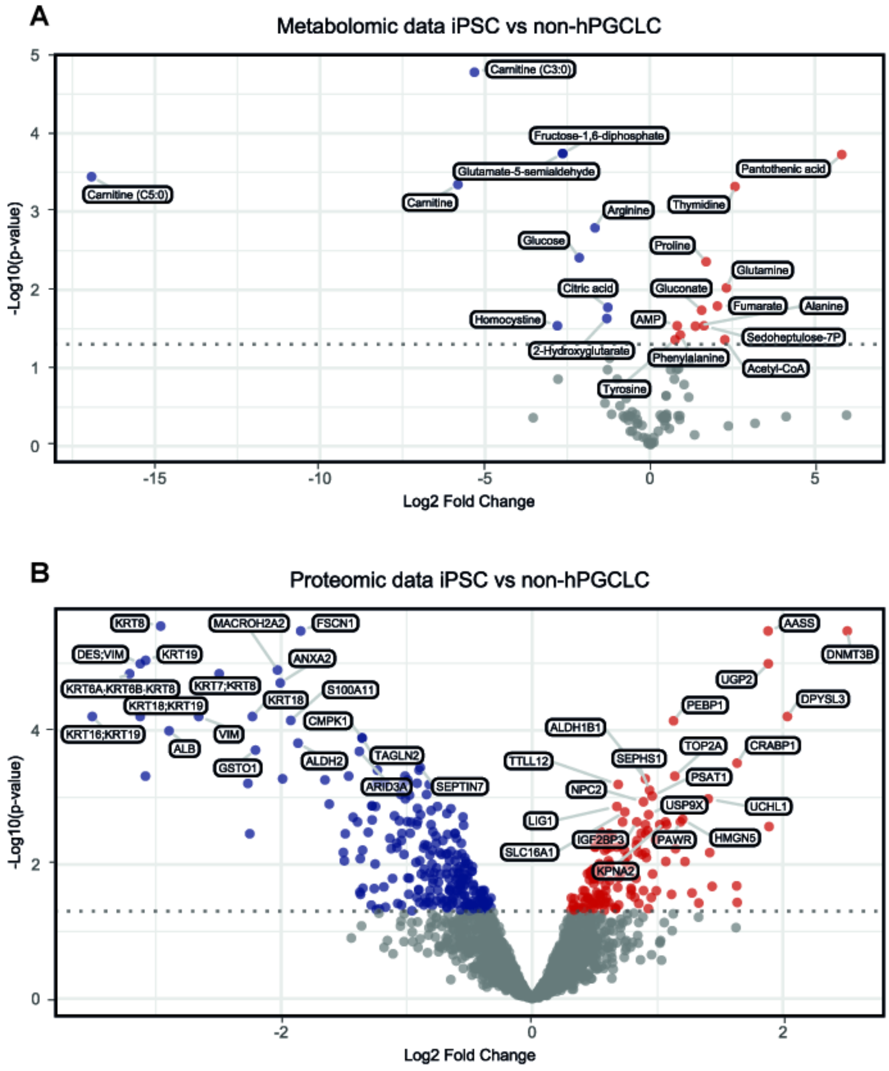


**Figure S2** – Metabolome and proteome comparison of iPSCs vs non-hPGCLCs, based on data from Table S5 and S6. **A**) Volcano plot of the metabolome comparison; metabolites organized by p-value and increased (red) or decreased (blue) abundance; **B**) Volcano plot of the proteome comparison; proteins organized by p-value and increased (red) or decreased (blue) expression.

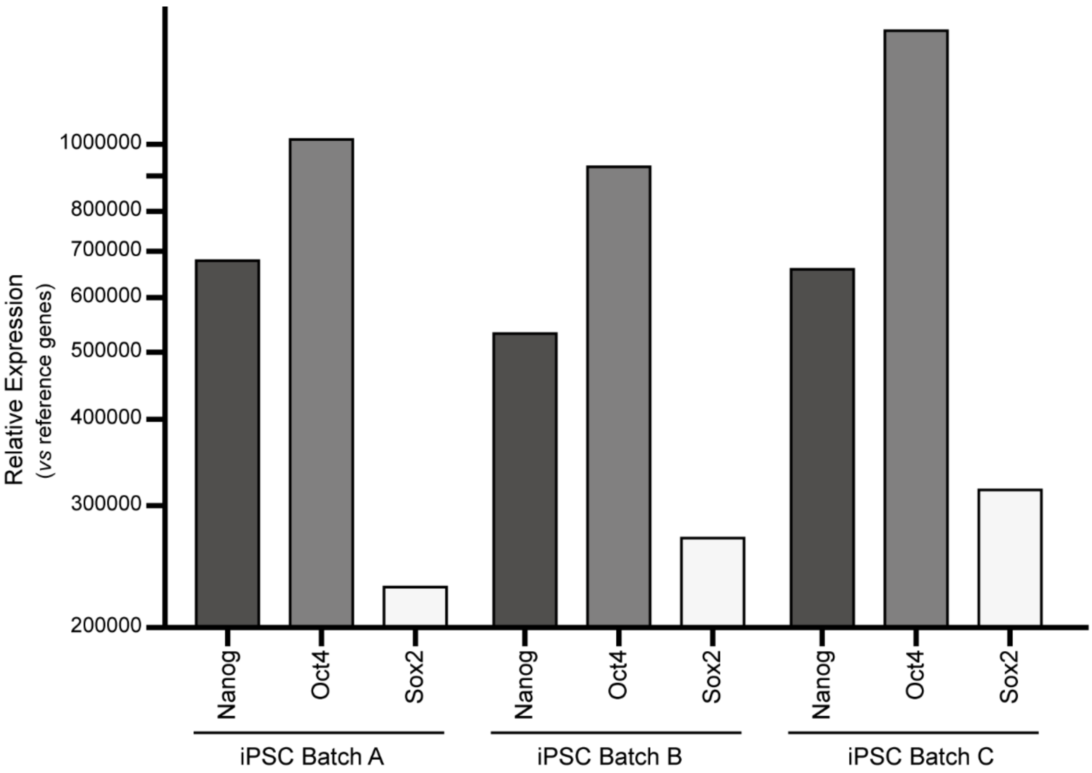


**Figure S3** – Validation of pluripotency marker's RNA expression in undifferentiated iPSCs. Quantitative PCR performed for *Nanog*, *Oct4* and *Sox2*. Use of three independent iPSC cultured samples, correspondent to distinct passage numbers: passage 23 (iPSC Batch A), passage 27 (iPSC Batch B) and passage 22 (iPSC Batch C). Relative Expression is based on Ncopy values of the target genes normalized against reference genes. Each bar corresponds to the geometric mean of three technical replicates (wells).
